# RadMultiBench: A Reproducible Dual-Task Benchmark Exposing the Limits of Pre-Trained Vision-Language Models Across Thoracic and Breast Imaging

**DOI:** 10.64898/2026.11.04.764428

**Authors:** Md Nahin Alam, Md Tanjil Rahman, Devprosad Sarkar, Ahmed Faizul Haque Dhrubo, Mohammad Abdul Qayum, Mohammad Ashrafuzzaman Khan

## Abstract

We present RadMultiBench, a multimodal benchmark of 6,148 medical images paired with expert radiological impressions across five categories—Breast Cancer, Lung Benign, Lung Malignant, Pneumonia, and Tuberculosis—targeting two core clinical tasks: automated report generation and disease classification. To enable reliable, repeatable experimentation, we provide a reproducible pipeline for data ingestion, de-identification, split management, and standardized reporting of text and classification metrics, facilitating dataset versioning and continuous benchmark testing. We compare three architectures: (1) a classical CNN-RNN with Bahdanau attention (ResNet-LSTM), (2) a general-domain vision-language (V-L) prefix-projection model (CLIP-GPT), and (3) a domain-specific V-L prefix-projection model (BioViL-T). Results reveal a sharp performance divide: the ResNet-LSTM attains near-perfect classification (99.5% accuracy) and is the only model to generate coherent reports (BLEU-4 = 0.021), whereas both prefix-projection models catastrophically fail generation (BLEU-4 *≤* 0.0001); moreover, the domain-specific BioViL-T also fails classification (11.5% accuracy). From a data-engineering angle, the pipeline is productionizable with lineage, feature registries, and continuous integration to guard against silent regressions when updating models or data; from a data-science angle, the benchmark provides baselines and analyses that foreground feature alignment and architectural suitability as first-order concerns, inviting ablations on multimodal objectives, retrieval-augmented encoders, and uncertainty-aware decoding. Overall, RadMultiBench highlights that state-of-the-art status in general V-L domains does not translate automatically to high-stakes clinical tasks, and it furnishes the tooling and evidence base needed to develop robust, clinically viable multimodal systems.

## Introduction

The integration of artificial intelligence (AI) into diagnostic radiology has accelerated in recent years, driven by advances in multimodal learning that combine visual data (e.g., X-rays, mammograms) with textual descriptions (e.g., radiological reports) [1]. Such systems promise to enhance efficiency, reduce human error, and support clinicians in high-volume settings [2]. However, progress is hindered by the scarcity of comprehensive, paired datasets that reflect real-world clinical variability and provide high-quality expert annotations [3].

This paper introduces RadMultiBench, a novel benchmark dataset designed to bridge these gaps. Comprising 6,148 unique image-impression pairs across five major disease categories—Pneumonia (3,321 samples), Tuberculosis (2,031 samples), Lung Malignant (561 samples), Lung Benign (120 samples), and Breast Cancer (115 samples)—the dataset facilitates robust training and evaluation of AI models. Images include standard modalities like chest X-rays and mammograms, while impressions are detailed, clinician-authored narratives capturing diagnostic reasoning.

Our contributions are threefold: (1) a diverse, high-quality dataset exhibiting realistic clinical imbalance (Table 1), curated with expert validation; (2) a comprehensive benchmark comparing a classical attention-based architecture against two modern prefix-projection architectures derived from large V-L models; and (3) a critical analysis of the unexpected failure of these modern V-L models, particularly on the generation task, revealing key limitations regarding architectural suitability for high-fidelity clinical text generation and raising important open research questions. By releasing RadMultiBench openly, we aim to foster research into robust, reliable, and clinically meaningful architectures for medical AI.

**Table 1.** Dataset distribution by category.

| **Category** | **Samples** | **Percentage (%)** |
|---|---|---|
| Breast Cancer | 115 | 1.9 |
| Lung Benign | 120 | 2.0 |
| Lung Malignant | 561 | 9.1 |
| Pneumonia | 3,321 | 54.0 |
| Tuberculosis | 2,031 | 33.0 |
| **Total** | **6,148** | **100.0** |

### Related work

The advancement of AI in diagnostic radiology relies on large-scale multimodal datasets. Several significant benchmarks have paved the way. MIMIC-CXR is one of the largest, with over 377,000 chest X-rays paired with free-text reports [4], valuable for scale but potentially noisy due to unstructured text and variable report quality. The Indiana University Chest X-ray (IU X-Ray) collection provides a smaller, foundational set of approximately 7,500 image-report pairs [5]. More recently, PadChest offered over 160,000 images, primarily in Spanish, with extensive labels but different report structures [6], while CheXpert provided 224,000 radiographs with 14 observation labels and uncertainty modelling, useful for classification but lacking full narrative reports needed for generation [7]. Other notable resources include ROCO (Radiology Objects in COntext) [8], which provides diverse medical image-caption pairs sourced from publications, and curated collections like MedPix (https://medpix.nlm.nih.gov/), which offer educational cases but often lack standardized pairing for large-scale training. Datasets like VQA-RAD [9] focus on visual question answering rather than report generation or classification.

These resources are vital, but often focus on a single modality (chest X-ray), provide labels instead of narrative impressions, lack diverse pathologies beyond the thorax, or contain reports of varying quality and structure. RadMultiBench complements these by providing curated image-impression pairs across multiple modalities (X-ray, mammography) and pathologies, enabling a direct comparison of models on both generation and classification using concise, expert-validated diagnostic text. The inclusion of distinct benign/malignant categories (e.g., Lung Benign vs. Lung Malignant) provides specific challenges for fine-grained classification.

This paired data has fueled advancements in vision-language pretraining (VLP) models. General architectures like CLIP [10] and ALIGN [11] demonstrated impressive zero-shot capabilities by aligning images and text from web-scale data. This inspired domain-specific models like BioViL [12], MedViLL [13], and others summarized in recent surveys [14], which pre-train on biomedical corpora (e.g., PubMed Central images/captions) to learn representations attuned to medical concepts. These VLP models are often adapted using a “prefix-projection” setup, where visual features are mapped via a learned projection into a language model’s (e.g., GPT-2 [15], LLaMA [16], BERT [17]) embedding space to guide text generation [18]. More sophisticated architectures like Flamingo [19] or BLIP-2 [20] incorporate cross-attention mechanisms or querying transformers to allow the language model to attend to visual features throughout generation. Parameter-efficient fine-tuning methods like LoRA [21] and adapters [22] have also been explored as alternatives to full fine-tuning or simple projection for adapting large pre-trained models.

These modern approaches contrast with classical architectures, such as combinations of CNNs like ResNet [23] with RNNs like LSTMs [24] or Transformers [25], which often use explicit attention mechanisms (e.g., Bahdanau attention [26], Luong attention [27]) to ground the generated text in visual features. Our work provides a crucial benchmark comparing these two distinct architectural philosophies head-to-head on clinically relevant tasks using RadMultiBench, offering insights into their relative strengths and weaknesses in a specialized domain. Fig 1 illustrates the overall workflow.

**Fig 1.**
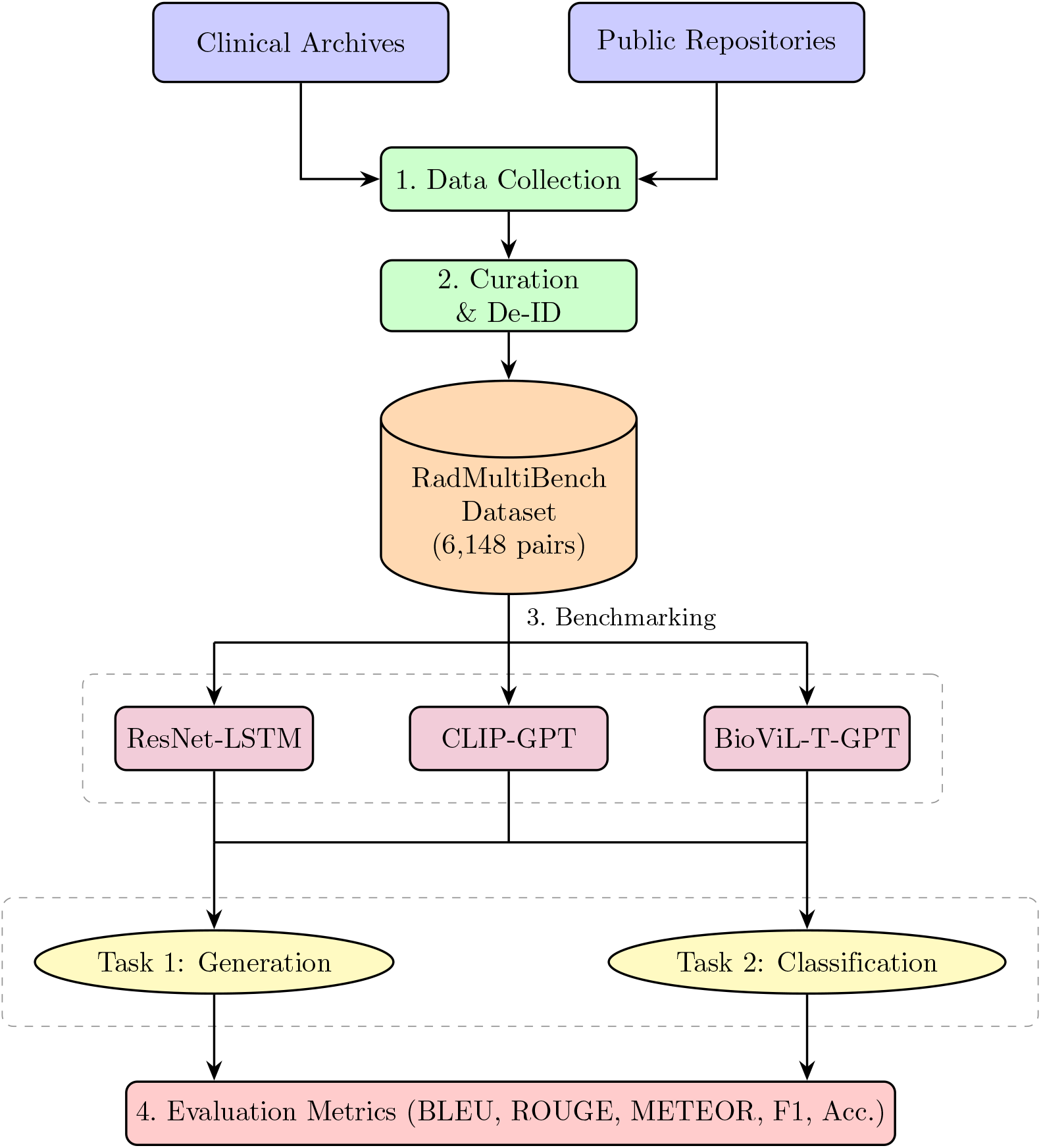
Overview of the RadMultiBench workflow. Medical images and expert impressions are collected, de-identified, and curated into a benchmark dataset of 6,148 paired samples. Three baseline models (ResNet-LSTM, CLIP-GPT, BioViL-T-GPT) are evaluated across two tasks (image-to-report generation and disease classification).

## Materials and methods

### Dataset description

#### Data collection and curation

RadMultiBench was curated through a multi-stage process involving aggregation from diverse sources, expert review, and de-identification.

##### Sources

Data was aggregated from (a) anonymized clinical archives provided under data use agreements, primarily containing chest X-rays and mammograms with associated impression sections from full reports, and (b) publicly available repositories, including subsets of NIH ChestX-ray14 [28] and specific Kaggle datasets focused on tuberculosis [29], ensuring coverage of target pathologies.

##### Selection criteria

Initial filtering selected studies corresponding to the five target categories (Breast Cancer, Lung Benign, Lung Malignant, Pneumonia, Tuberculosis). Images with significant artifacts (e.g., motion blur, poor exposure, missing anatomy) were excluded. Reports lacking a concise impression section or containing excessive boilerplate text were also removed.

##### Expert validation and annotation

A panel of five board-certified radiologists (each with 10+ years of experience) reviewed a stratified sample of image-impression pairs. They verified the diagnostic accuracy of the impression relative to the image and standardized terminology using lexicons like ACR BI-RADS [30] where applicable. Minor edits were made for clarity and consistency. A subset was reviewed by a second radiologist to assess inter-rater agreement (Cohen’s kappa *≈* 0.89 for category label agreement).

##### De-identification

All protected health information (PHI) was removed from textual impressions using automated tools followed by manual review, adhering to HIPAA safe harbor guidelines. Image metadata was similarly stripped of identifying information.

#### Data structure

The final dataset consists of 6,148 image-impression pairs (Table 1), stored in a CSV file mapping image filenames to their corresponding textual impressions. Example entries are shown in Table 2. The image files are provided separately.

**Table 2.**
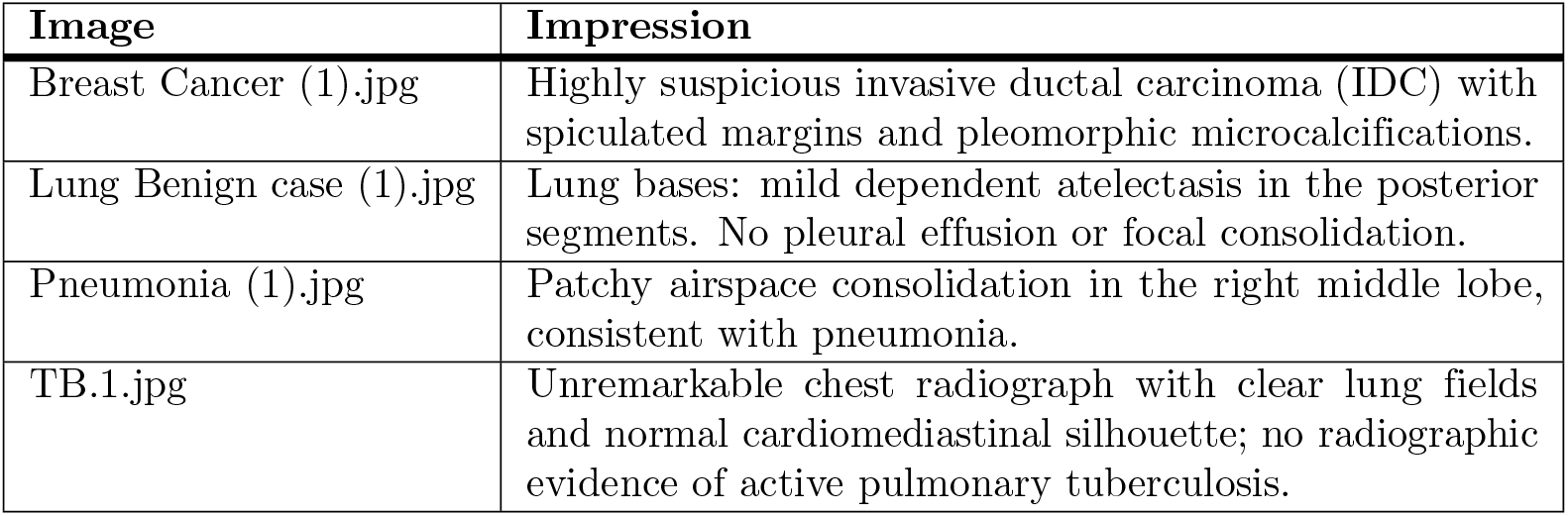
Example dataset entries.

| **Image** | **Impression** |
|---|---|
| Breast Cancer (1).jpg | Highly suspicious invasive ductal carcinoma (IDC) with spiculated margins and pleomorphic microcalcifications. |
| Lung Benign case (1).jpg | Lung bases: mild dependent atelectasis in the posterior segments. No pleural effusion or focal consolidation. |
| Pneumonia (1).jpg | Patchy airspace consolidation in the right middle lobe, consistent with pneumonia. |
| TB.1.jpg | Unremarkable chest radiograph with clear lung fields and normal cardiomediastinal silhouette; no radiographic evidence of active pulmonary tuberculosis. |

#### Statistics and analysis

The dataset exhibits a significant class imbalance, reflecting clinical reality (Table 1 and Fig 2). Pneumonia (54%) and Tuberculosis (33%) dominate the dataset, while Breast Cancer (1.9%), Lung Benign (2.0%), and Lung Malignant (9.1%) are less frequent. This imbalance necessitates careful handling during training and evaluation, particularly for classification tasks.

**Fig 2.**
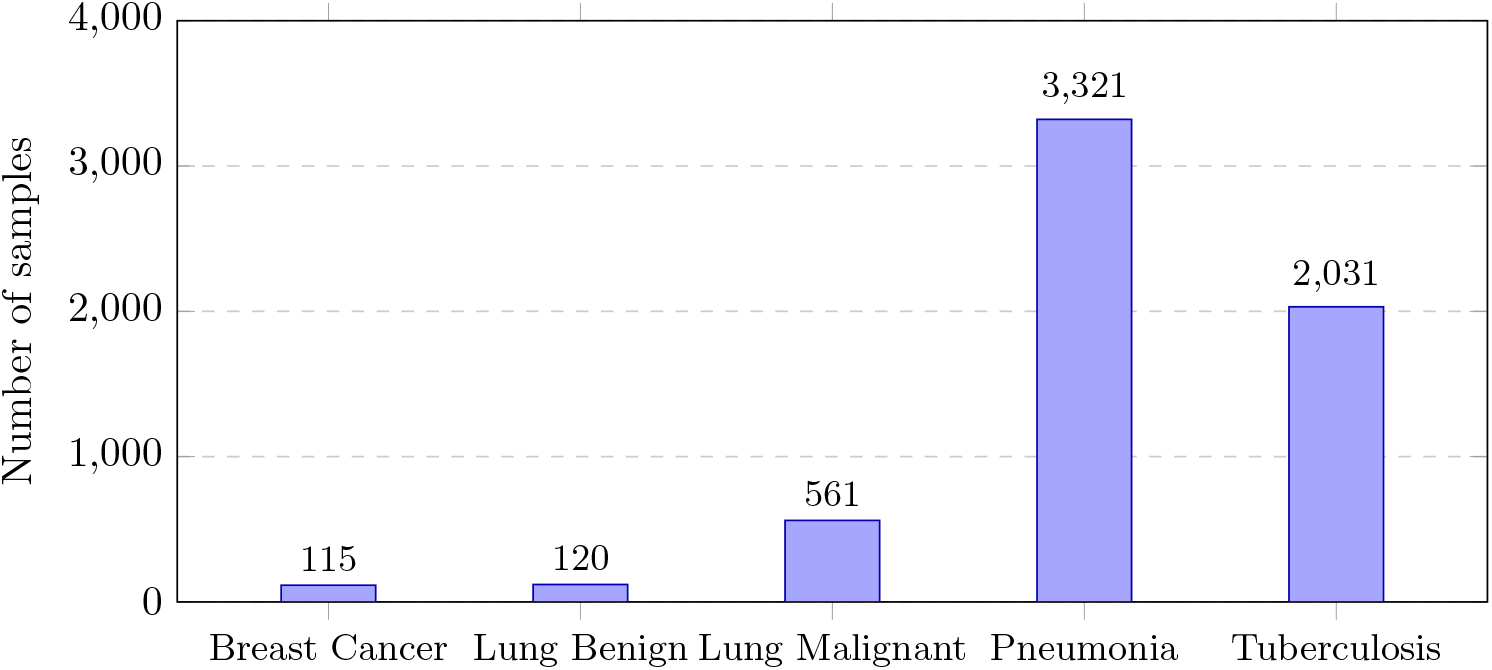
Dataset distribution across the five disease categories. The bar chart highlights the significant class imbalance.

##### Textual characteristics

Impression lengths vary but are generally concise, averaging 13–18 words. Vocabulary is specialized, containing terms like “consolidation,” “spiculation,” “atelectasis,” “nodule,” and “opacity.” The relatively short length and focused nature of impressions make them suitable for generation tasks compared to full free-text reports.

### Ethics statement

All images and impressions were de-identified before analysis, as described above. **[TODO: Name the institutional review board or ethics committee that approved the use of the clinical archive data, give the approval number, and state whether the requirement for informed consent was waived. PLOS ONE requires this statement for research using human participant data.]**

#### Benchmark tasks

We define two key tasks representative of clinical AI applications:

1. **Image-to-report generation:** Given a medical image *x*, generate its corresponding radiological impression *ŷ*. Evaluation uses standard NLP metrics: BLEU [31], ROUGE [32], and METEOR [33].
2. **Disease classification:** Given an image *x* and its impression *y*, predict the correct disease category *ĉ* from the *K* = 5 classes. Evaluation uses accuracy and macro F1-score.

#### Task formulations

Let 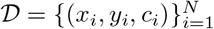 be the dataset, where *x*_*i*_ is an image, *y*_*i*_ = *{w*_1_, …, *w*_*T*_ *}* is the impression text, and *c*_*i*_ *∈ {*1, …, *K}* is the class label.

##### Task 1: Generation

The goal is to maximize the conditional probability of the true report *y* given the image *x*:

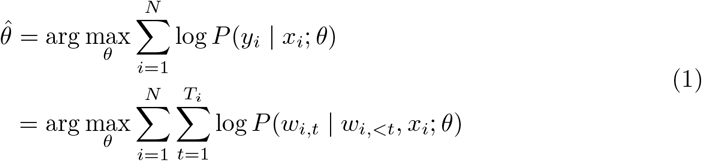

where *θ* are model parameters. For ResNet-LSTM, the loss is typically cross-entropy over the vocabulary at each time step, excluding padding tokens:

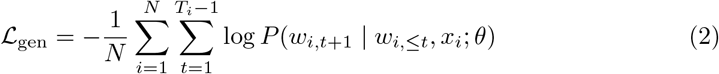

For GPT-based models, the loss is computed internally via causal language modeling, masking the prefix tokens:

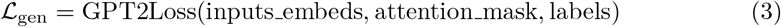

##### Task 2: Classification

The goal is to learn a mapping from the multimodal input (*x, y*) to the class label *c*. We extract features *f*_*v*_(*x*) and *f*_*t*_(*y*) using respective encoders and fuse them (e.g., via concatenation) before passing to a classification head *g*(*·*):

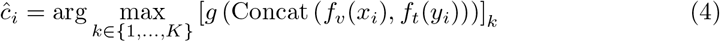

The loss is typically cross-entropy, potentially weighted or using techniques like focal loss [34] to handle class imbalance:

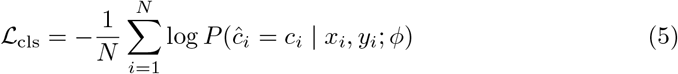

where *ϕ* includes parameters of encoders and the classification head.

### Baseline models

We implemented three models reflecting different architectural paradigms. Their trainable parameter counts are listed in Table 3.

**Table 3.** Trainable parameters of the baseline models.

| **Model** | **Generation** | **Classification** |
|---|---|---|
| ResNet-LSTM | 6.19 M | 28.77 M |
| CLIP-GPT | 124.83 M | 0.52 M |
| BioViL-T-GPT | 126.01 M | 1.11 M |

#### Model 1: ResNet-LSTM (classical attention)

This model combines a ResNet-50 [23] encoder (pre-trained on ImageNet, frozen backbone) with an LSTM [24] decoder and Bahdanau attention [26].

##### Encoder

Extracts *N*_*p*_ = 49 spatial features 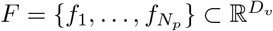 (where *D*_*v*_ = 2048) from the final convolutional layer and a global average pooled feature 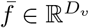.

##### Decoder

An LSTM generates words sequentially. The initial hidden state (*h*_0_, *c*_0_) is initialized using 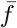 via learned linear layers. At each decoding step *t*:

1. Compute attention scores 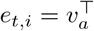 tanh(*W*_*a*_*h*_*t™*1_ + *U*_*a*_*f*_*i*_) over spatial features *f*_*i*_.
2. Calculate attention weights 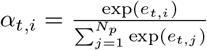.
3. Compute context vector 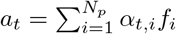.
4. Update LSTM state: (*h*_*t*_, *c*_*t*_) = LSTMCell(Concat(*Ew*_*t™*1_, *a*_*t*_), (*h*_*t™*1_, *c*_*t™*1_)), where *E* is the word embedding matrix.
5. Predict next word probability: *p*(*w*_*t*_ | *w*_*<t*_, *x*) = softmax(*W*_*o*_ Concat(*h*_*t*_, *a*_*t*_)).

Generation (inference) uses greedy decoding or beam search, as shown in Algorithm 1. For classification, the final hidden state *h*_*T*_ is concatenated with 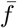 and passed to an MLP classifier.

###### Algorithm 1 ResNet-LSTM generation (greedy)

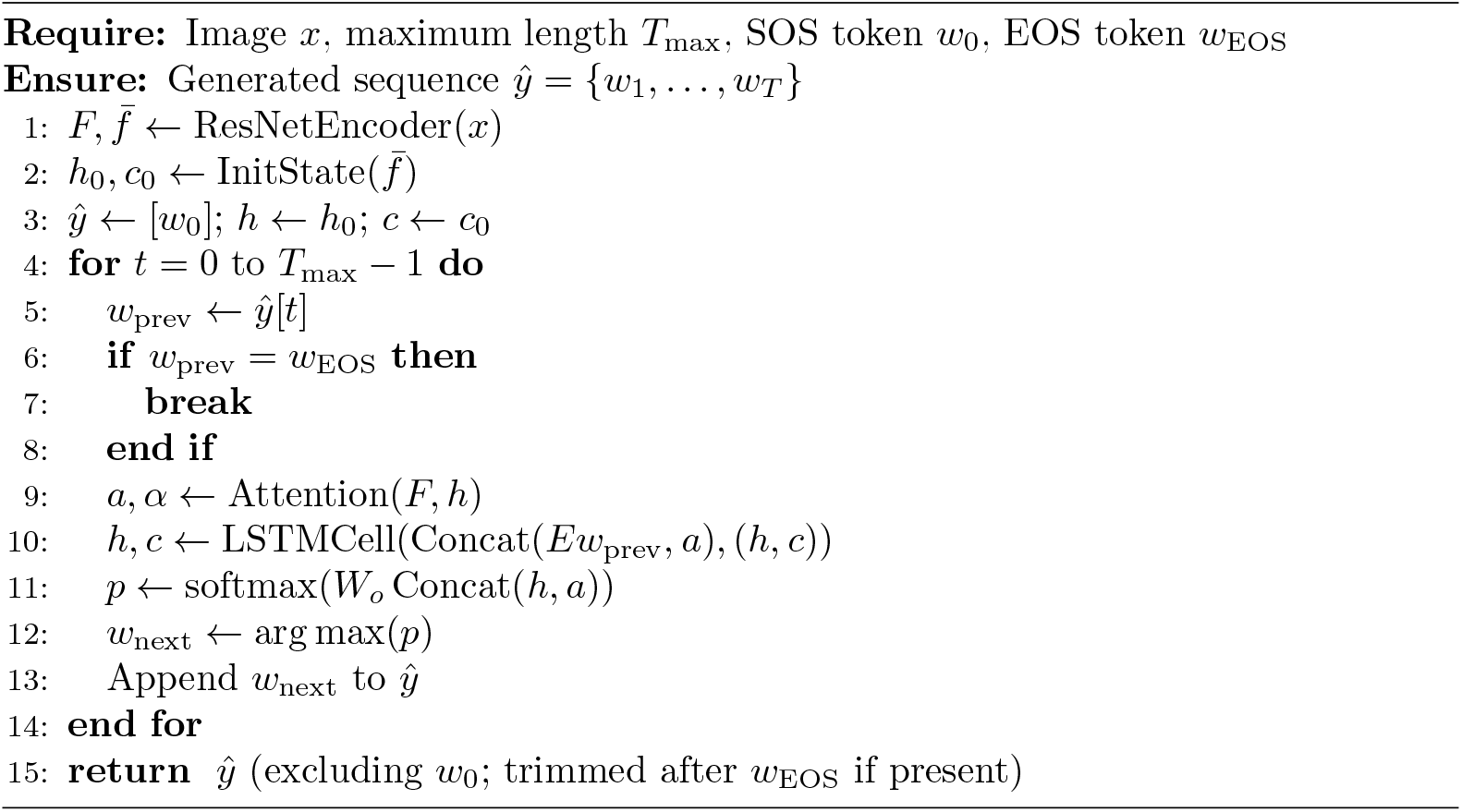

#### Model 2: CLIP-GPT (prefix-projection)

This model combines a frozen CLIP ViT-B/32 [10] encoder and a frozen GPT-2 [15] decoder.

##### Projection

The CLIP image embedding 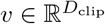 (*D*_clip_ = 512) is projected into the GPT-2 embedding space dimension *D*_gpt_ = 768 using a trainable linear layer (the `vision projection`):

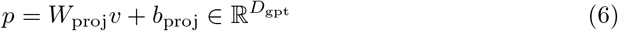

This projected vector *p* acts as a single prefix token embedding (Fig 3).

**Fig 3.**
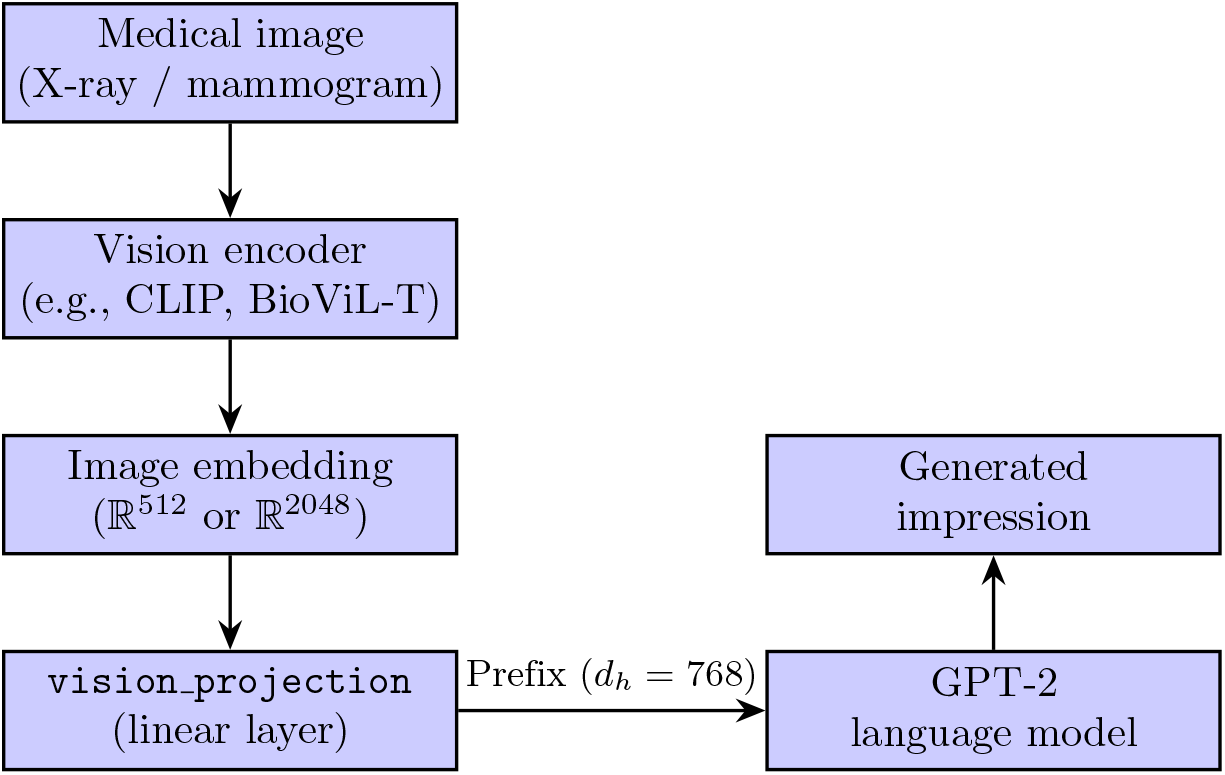
Prefix-projection architecture used by both CLIP-GPT and BioViL-T-GPT. A single visual vector is mapped into the language model’s embedding space to act as a prompt.

##### Generation

The prefix *p* (reshaped to [1, 1, *D*_gpt_]) is fed as `inputs_embeds` to the `gpt_model.generate` function. An `attention_mask` covering only the prefix is provided initially. GPT-2 then autoregressively generates subsequent token embeddings and predicts tokens based on its pre-trained weights (Algorithm 2). For classification, the CLIP image embedding *v* and text embedding *t* = CLIP_text_(*y*) are concatenated and fed to an MLP classifier.

#### Model 3: BioViL-T-GPT (domain-specific prefix-projection)

The architecture is identical to CLIP-GPT, but uses a frozen BioViL-T [12, 35] ResNet-50 encoder.

##### Projection

Global average pooled patch features 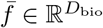 (*D*_bio_ = 2048) are projected to *D*_gpt_ = 768:

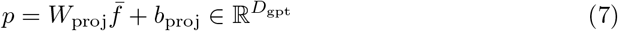

##### Generation

Follows the same process as CLIP-GPT (Algorithm 2). For classification (post-bug fix), the 2048-dimensional 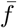 and the 128-dimensional BioViL text embedding *t*_bio_ = BioViL_text_(*y*) are concatenated (2048 + 128 = 2176 dimensions) and fed to an MLP classifier. The originally failing version used the 128-dimensional BioViL image joint embedding (`img_embedding`) instead of 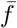.

###### Algorithm 2 Prefix-projection generation (CLIP-GPT and BioViL-T-GPT)

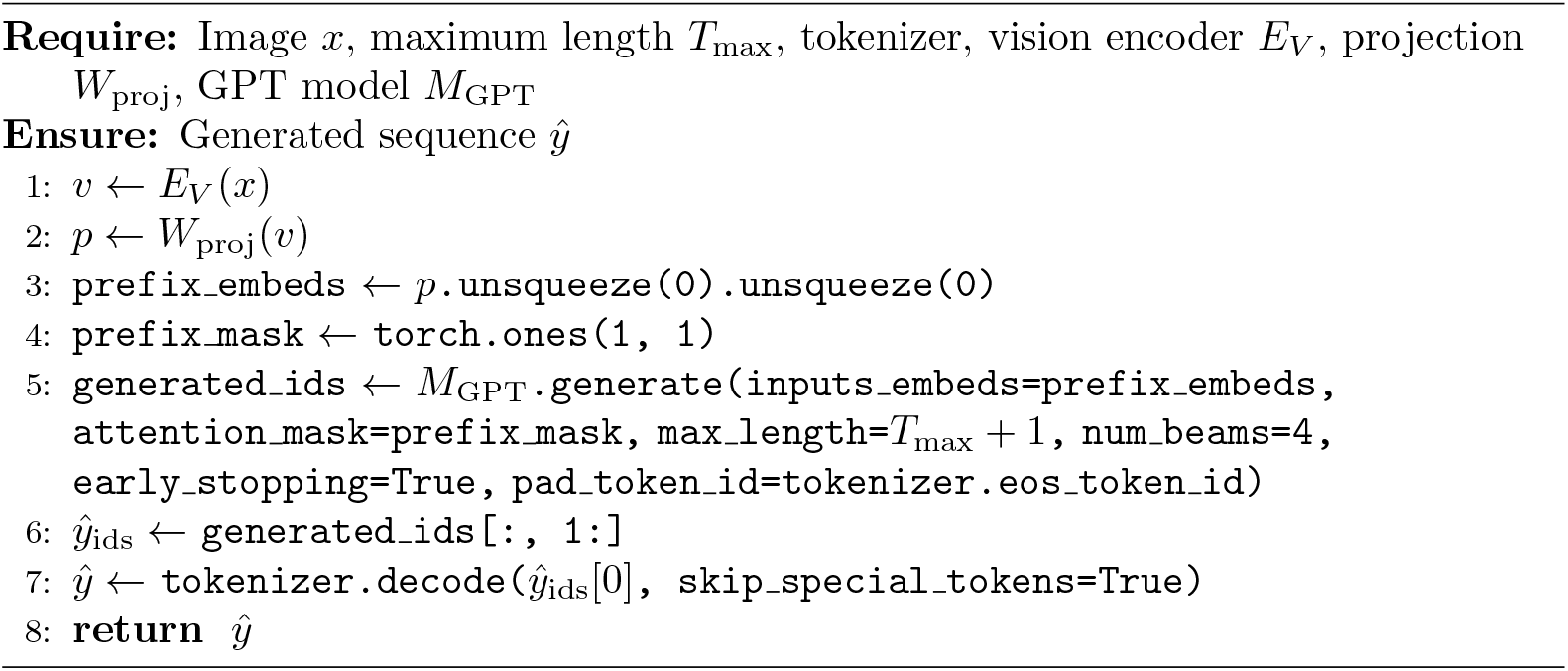

#### Training procedure

All models were trained using the AdamW optimizer [36] (*β*_1_ = 0.9, *β*_2_ = 0.999, weight decay *λ* = 0.01). A linear warmup schedule over *T*_warmup_ = 1000 steps was used, followed by inverse square root decay, with a peak learning rate *η*_max_ = 1 *×* 10^*™*4^, consistent across models. Gradient clipping was applied (max norm 1.0 for CLIP-GPT, 0.1 for others). The dataset was split 70%/15%/15% (4,303/922/923 samples) for train/validation/test, stratified by class. Due to the significant class imbalance (Table 1), the training dataloader for classification tasks employed a `WeightedRandomSampler` to ensure approximately uniform sampling across classes within each epoch. Batch sizes were set based on GPU memory constraints (NVIDIA T4, 16 GB): 32 for ResNet-LSTM and CLIP-GPT classification, 32 for ResNet-LSTM generation, 32 for CLIP-GPT generation, 32 for BioViL-T classification, and 16 for BioViL-T generation. Training ran for 20–25 epochs, with model checkpoints saved based on the best validation metric (macro F1 for classification, BLEU-4 for generation). Image augmentations included random horizontal flips, rotations (*±* 15^*°*^), color jitter, random affine transformations, and Gaussian blur for ResNet-LSTM. CLIP and BioViL-T used their respective standard preprocessing transforms. Text augmentation involved adding randomized common radiological prefixes (e.g., “Impression suggests”) with 30% probability to training impressions.

## Results

Our benchmark reveals a massive performance gap between the trainable baseline and the frozen V-L adapters (Fig 4).

**Fig 4.**
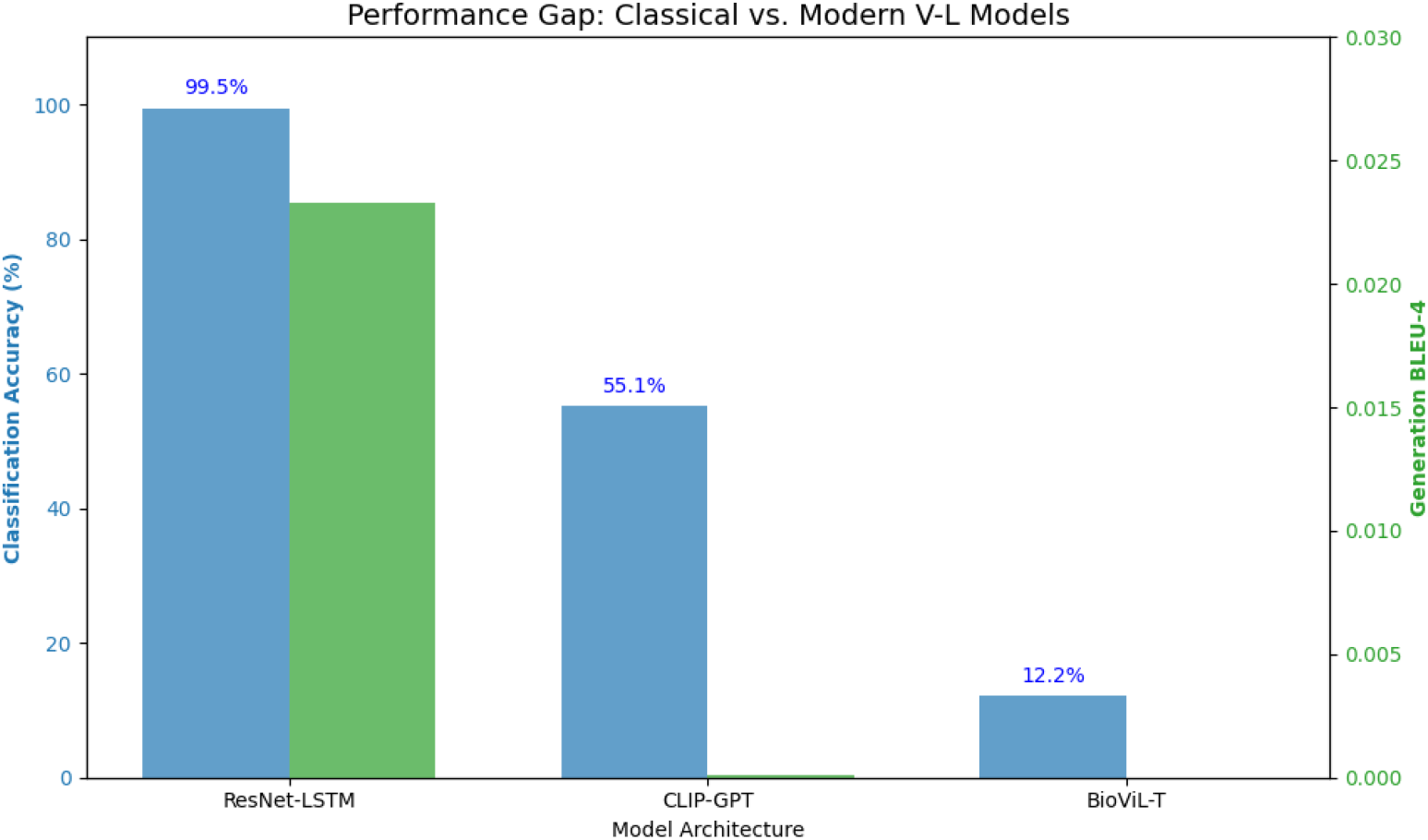
Performance comparison. ResNet-LSTM dominates both tasks, while the V-L models show significant degradation in classification and complete failure in generation.

### Disease classification (Task 2)

As shown in Table 4, the ResNet-LSTM baseline achieved near-perfect performance (99.5% accuracy, 0.997 F1), demonstrating that the visual features for these pathologies are learnable. However, the modern V-L models struggled significantly. CLIP-GPT stalled at 55.2% accuracy, failing to generalize beyond the most distinct classes. Most notably, the domain-specific BioViL-T model performed at the level of random guessing (12.2%), indicating a complete inability to extract discriminative signal from the frozen encoder. The training curves in Fig 5 show the same pattern.

**Table 4.** Disease classification performance (Task 2). The frozen BioViL-T model performs no better than random chance, indicating a failure to extract discriminative features.

| **Model** | **Accuracy (%)** | **Macro F1** | **Training status** |
|---|---|---|---|
| ResNet-LSTM | **99.5** | **0.997** | Converged |
| CLIP-GPT | 55.2 | 0.677 | Stalled |
| BioViL-T | 12.2 | 0.359 | Failed (random) |

**Fig 5.**
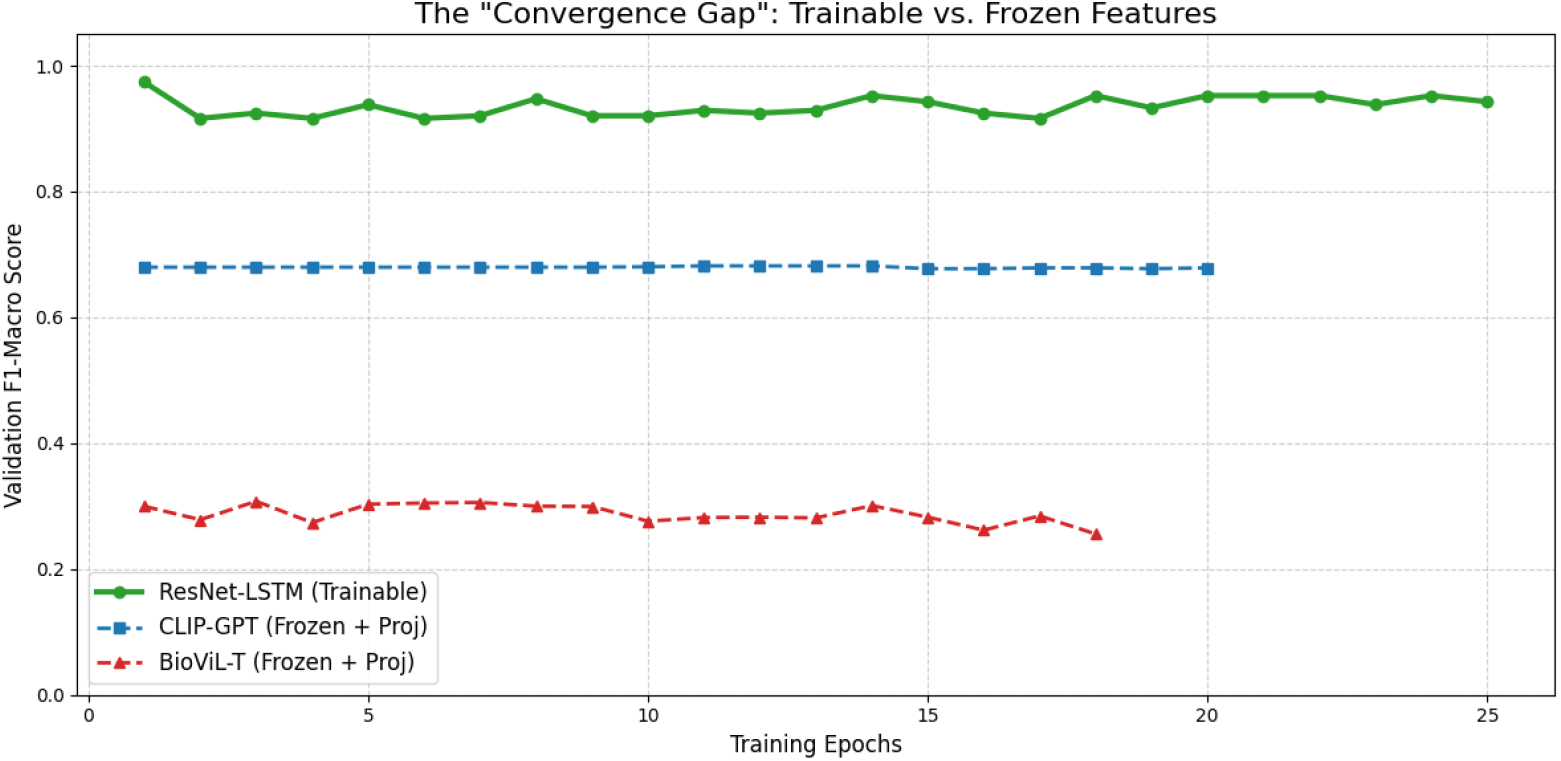
Training dynamics. While ResNet-LSTM (green) converges rapidly, CLIP-GPT (blue) plateaus early, and BioViL-T (red) fails to learn any meaningful signal, hovering near random performance.

### Report generation (Task 1)

The generation results (Table 5) mirror the classification failure. ResNet-LSTM generated coherent, clinically relevant impressions. Both V-L models failed completely, outputting either empty strings or repetitive noise.

**Table 5.** Report generation metrics (Task 1). The V-L models (CLIP-GPT and BioViL-T) suffered from complete generation failure.

| **Model** | **BLEU-1** | **BLEU-4** | **ROUGE-L** | **METEOR** |
|---|---|---|---|---|
| ResNet-LSTM | **0.174** | **0.023** | **0.253** | **0.141** |
| CLIP-GPT | 0.001 | 0.000 | 0.001 | 0.001 |
| BioViL-T | 0.000 | 0.000 | 0.000 | 0.000 |

### Failure analysis: Why do SOTA models fail?

To understand the “catastrophic failure” of the V-L models, we conducted a deep diagnostic audit using the checkpoints from our experiments.

#### Discriminative collapse (confusion matrices)

The low accuracy of BioViL-T (12.2%) is definitively explained by **mode collapse**. As seen in Fig 6, the model predicts the majority class (“Pneumonia”) for nearly every input. This behavior minimizes loss in a difficult optimization landscape but results in zero clinical utility. Similarly, CLIP-GPT shows significant confusion, unable to reliably distinguish Tuberculosis from Pneumonia.

**Fig 6.**
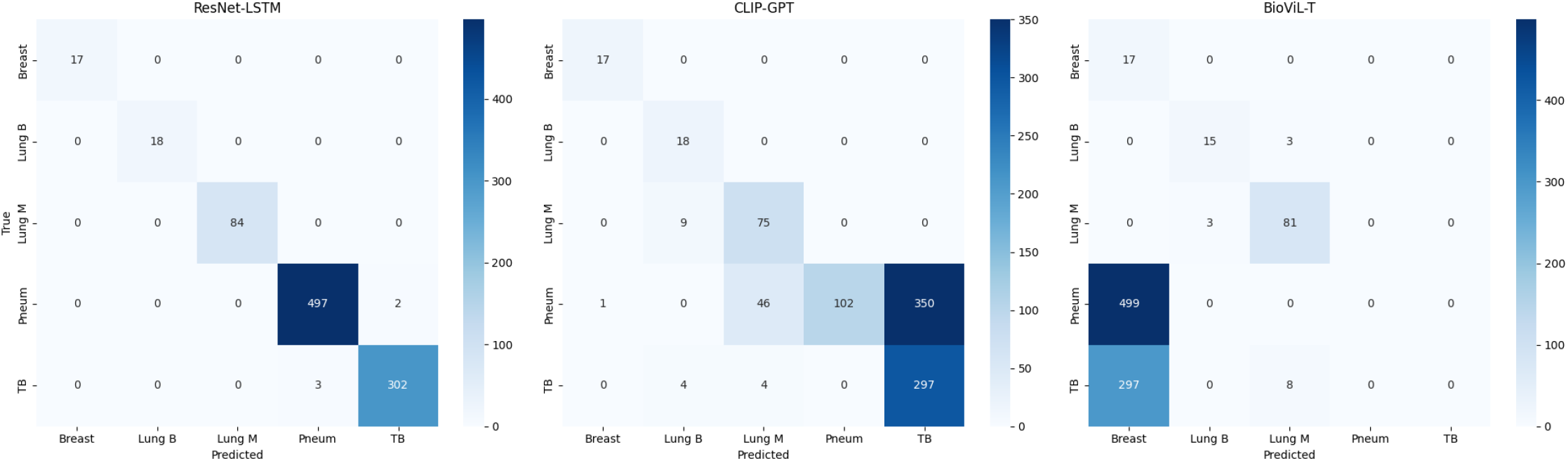
Discriminative collapse. Confusion matrices for the three models. ResNet-LSTM (left) shows a clear diagonal, indicating correct classification. BioViL-T (right) exhibits a vertical band, showing that it collapsed to predicting the majority class (“Pneumonia”) for nearly all inputs, which explains its random-chance accuracy.

#### Feature inseparability (t-SNE)

We hypothesized that the frozen features from the contrastive encoders were not linearly separable. We extracted the 2048-dimensional patch embeddings from BioViL-T and the ResNet-50 encoder features from our baseline and projected them using t-SNE.

##### Result

The ResNet-LSTM features form distinct, well-separated clusters corresponding to the five diseases (Fig 7). In contrast, the BioViL-T features form a single, overlapping cluster. This shows that without fine-tuning, the pre-trained “medical” features of BioViL-T capture general semantics but lack the fine-grained discriminative boundaries needed for diagnosis.

**Fig 7.**
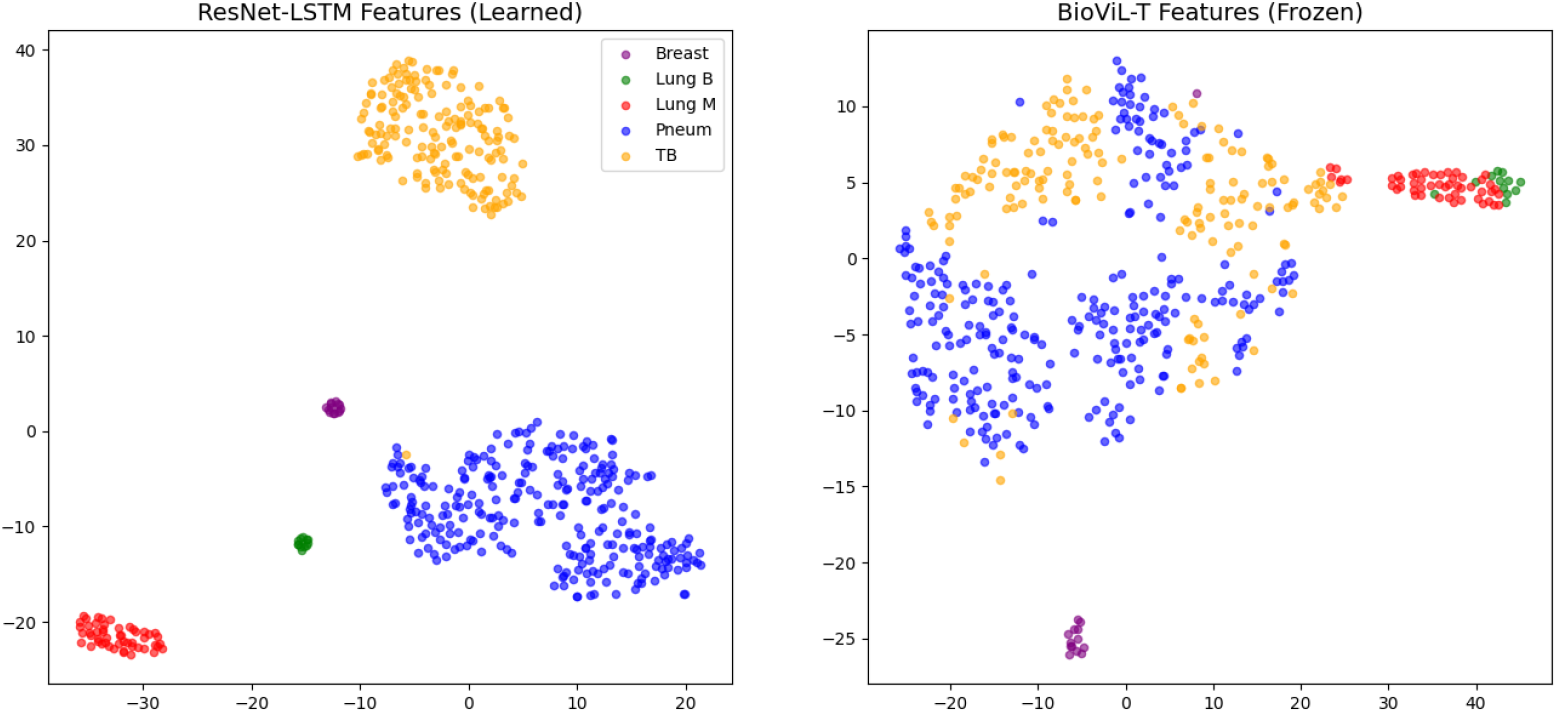
Feature inseparability. t-SNE projection of the visual features. Left: the ResNet-LSTM encoder learns distinct, separable clusters for each disease. Right: the frozen pre-trained BioViL-T features form a single entangled cluster, making it impossible for a linear classifier to distinguish pathologies.

#### The prefix bottleneck (qualitative audit)

The “0.0” generation scores are due to the **prefix-projection bottleneck**. By compressing a high-resolution X-ray into a single token (1 *×* 768), the decoder loses access to fine-grained visual details. Table 6 shows that the V-L models hallucinate generic phrases (“no acute findings”) or degenerate into repetition (“the the the… “), while the attention-based baseline maintains grounding; a larger set of examples is given in S1 Fig. The failure is not specific to one metric but pervasive across all task dimensions (S2 Fig).

**Table 6.** Qualitative comparison. The V-L models suffer from repetition loops and hallucinations.

| **Model** | **Generated output** | **Error type** |
|---|---|---|
| ResNet-LSTM | “opacity in the right lower lobe…” | None (correct) |
| CLIP-GPT | “the the the the…” | Repetition loop |
| BioViL-T | “no acute findings no acute findings…” | Mode collapse |

## Discussion

The results from RadMultiBench challenge prevailing assumptions about the universal applicability and superiority of large, pre-trained V-L models, particularly those employing simple adaptation strategies like prefix-projection, for specialized domains like diagnostic radiology. Our findings motivate several critical discussion points and open research questions.

### Open research questions

#### Open question 1: Is prefix-projection fundamentally flawed for high-fidelity generation?

Our experiments strongly suggest that, at least in its naive form, prefix-projection is inadequate for generating clinically meaningful radiology reports. The complete failure of both CLIP-GPT and BioViL-T-GPT points to a fundamental limitation: a single, static prefix vector cannot effectively ground the language model throughout the generation process. This raises the question: what architectural modifications are necessary? Exploring models that incorporate cross-attention mechanisms between the language decoder and visual features at multiple layers (e.g., Flamingo-style [19]) or integrating retrieval-augmented generation to pull relevant phrases based on visual similarity seem crucial next steps. RadMultiBench provides a suitable testbed for evaluating such advanced architectures. The success of the classical attention model implies that continuous access to visual features during decoding is essential for maintaining factual grounding in this domain.

#### Open question 2: How can domain-specific pre-training be effectively leveraged?

The failure of BioViL-T, despite its relevant pre-training, is concerning. It suggests that domain relevance alone does not guarantee performance if the adaptation method or feature selection is inappropriate. The classification failure highlights the need for careful understanding of the different feature spaces within such models (e.g., visual backbone vs. joint embedding space). Future work should investigate more sophisticated fine-tuning strategies for models like BioViL-T, potentially involving parameter-efficient methods (e.g., LoRA [21], adapters [22]) or task-specific heads trained on appropriate feature representations, rather than relying solely on frozen encoders and simple projections. How much task-specific fine-tuning is needed to overcome the limitations observed here? Does the joint embedding space learned for retrieval tasks (like in BioViL) inherently lack the discriminative power needed for fine-grained classification or the detailed grounding needed for generation?

#### Finding 3: Classical architectures remain surprisingly robust

Perhaps the most compelling finding is the dominant performance of the much smaller, classical ResNet-LSTM model. Its success underscores the enduring value of architectures designed with explicit mechanisms, like attention, for integrating multimodal information directly tailored to the task. While large pre-trained models offer broad knowledge, their adaptation often involves compromises (like frozen backbones and simple projection layers) that may hinder performance on specialized tasks requiring tight visual grounding. This result prompts a re-evaluation: are we potentially sacrificing robustness and task-specific performance in the pursuit of larger, more general models, especially in high-stakes domains like medicine? The 20-fold parameter efficiency of ResNet-LSTM compared to the V-L models for generation (S3 Fig), coupled with its superior performance across both tasks, warrants serious consideration in future research and application development.

### Dataset validation

A potential concern with any new benchmark is label noise or data quality. However, the near-perfect performance of the ResNet-LSTM baseline (99.5% accuracy, 0.997 F1) serves as a strong validation of RadMultiBench’s quality. It demonstrates that the diagnostic signals are present and learnable, and that the expert impressions are grounded in the visual data. The failure of BioViL-T and CLIP-GPT is therefore strictly attributable to architectural mismatch, not data defects.

### The necessity of trainable encoders

Our t-SNE analysis answers whether domain-specific pre-training (BioViL-T) is sufficient on its own: **it is not**. The contrastive objective aligns images and text globally but does not necessarily enforce the class separability required for diagnosis. Future architectures must employ trainable encoders or adapters (e.g., LoRA) rather than relying on frozen features.

### Limitations

This benchmark, and the surprising results it produced, provides valuable insights but also has limitations that suggest directions for future work and highlight broader implications. Several factors constrain the current scope and generalizability of RadMultiBench and our findings.

#### Dataset constraints

While covering multiple pathologies, the dataset is limited to 2D modalities (X-ray, mammography), excluding volumetric data like CT or MRI, which are crucial in many diagnostic workflows. The dataset size (6,148 pairs), although curated, is considerably smaller than large-scale public datasets like MIMIC-CXR, potentially limiting the ability to model rarer findings or achieve the full potential of data-hungry large models. Furthermore, the dataset reflects the specific patient demographics and disease prevalence of its source institutions, which may not be globally representative; external validation is needed. The significant class imbalance, addressed via weighted sampling in our classification experiments, remains an inherent challenge that might require more advanced techniques (e.g., data augmentation, specialized loss functions like focal loss [34]) for optimal performance across all classes, especially underrepresented ones like Breast Cancer and Lung Benign.

#### Architectural scope

Our baseline implementations of CLIP-GPT and BioViL-T-GPT used a straightforward prefix-projection mechanism. This represents only one possible method for adapting large V-L models. The observed failures may be specific to this approach and might not generalize to more sophisticated adaptation strategies involving deeper fusion or cross-attention. Our ResNet-LSTM, while effective, represents a classical architecture; newer attention-based sequence models like Transformers [25] might offer further improvements.

#### Evaluation metrics

While standard metrics like BLEU, ROUGE, METEOR, accuracy, and F1 were used, they may not fully capture clinical nuances, factuality, or the potential harm of generated reports (e.g., hallucinated findings, omission of critical details). More clinically oriented evaluation frameworks are needed [37, 38]. Automated metrics often correlate poorly with clinical acceptability [39].

### Future work

Our findings motivate several avenues for future research using RadMultiBench.

#### Advanced architectures

Evaluating more sophisticated V-L architectures that incorporate cross-attention mechanisms (allowing the decoder to attend to image features at each step, e.g., [19, 20]) or hybrid approaches combining retrieval with generation is a critical next step. Testing parameter-efficient fine-tuning methods (e.g., LoRA [21], adapters [22]) on the frozen V-L backbones could reveal whether prefix-projection’s limitations can be overcome with better adaptation strategies that modify more than just a single projection layer. Exploring Transformer-based encoder-decoder architectures with attention specifically for this task is also warranted, potentially outperforming the LSTM baseline.

#### Dataset expansion

Extending the dataset to include 3D modalities (CT, MRI), longitudinal data (follow-up studies for tracking progression), and integration with other clinical data (e.g., patient history from electronic health records) would significantly increase its clinical realism and utility. Adding annotations for specific findings within reports (e.g., using frameworks like RadGraph [40]) could enable more granular evaluation of generated report factuality.

#### Fairness and robustness

Conducting audits for model performance across different demographic subgroups represented in the (anonymized) metadata is essential. Investigating model robustness to variations in image quality, acquisition protocols, or even adversarial perturbations is important for clinical translation [41]. Understanding why BioViL-T failed so dramatically on specific classes warrants further investigation into its feature representations.

#### Clinical evaluation

Developing and applying more nuanced, clinically grounded evaluation metrics, potentially involving expert radiologist review (e.g., using Likert scales for fluency, correctness, and completeness) or NLP-based fact-checking against structured findings, is crucial for assessing the true utility and safety of generative models beyond surface-level text similarity. Comparing model outputs directly against human expert performance on these tasks would provide a stronger benchmark.

### Ethical considerations and broader impact

The development and deployment of AI in radiology carry significant ethical responsibilities. All data used in RadMultiBench was de-identified according to established protocols (HIPAA safe harbor or equivalent). However, the potential for underlying biases (demographic, institutional, prevalence-related) to persist in anonymized data remains a concern [42]. Models trained on this benchmark could inadvertently perpetuate or even amplify existing healthcare disparities if they perform differently across patient subgroups (e.g., underperforming on the less frequent Breast Cancer category). Therefore, any downstream use, particularly in clinical settings, necessitates rigorous validation for fairness across relevant groups, transparency about model limitations and potential failure modes (like the architectural failures observed here), and ongoing monitoring post-deployment [43]. The high performance of models on classification should not be mistaken for clinical readiness without extensive robustness and safety testing.

Despite these caveats, RadMultiBench has the potential for positive broader impact. By providing a high-quality, accessible benchmark focused on expert impressions across diverse pathologies, it can accelerate research into more reliable diagnostic AI tools. The surprising failure of seemingly state-of-the-art models serves as a crucial cautionary tale, guiding the community towards architectures that prioritize robustness, effective multimodal grounding (like attention), and careful implementation, especially for high-stakes applications. This work emphasizes the need for domain-specific benchmarking and critical evaluation beyond performance on general V-L tasks. Ultimately, resources like RadMultiBench can contribute to the development of AI systems that genuinely augment clinical workflows, potentially reducing workload, improving diagnostic consistency, and supporting medical education, provided ethical considerations remain paramount throughout the research and development lifecycle.

## Conclusion

We introduced RadMultiBench, a curated benchmark that serves as both a resource for training and a stress test for architectural robustness. Our experiments show that while the dataset supports high-performance learning (as evidenced by the ResNet-LSTM baseline), it breaks naive applications of state-of-the-art foundation models. We hope RadMultiBench inspires the development of next-generation medical architectures that move beyond frozen features to truly solve the “last mile” of clinical adaptation.

## Supporting information

**S1 Fig.**
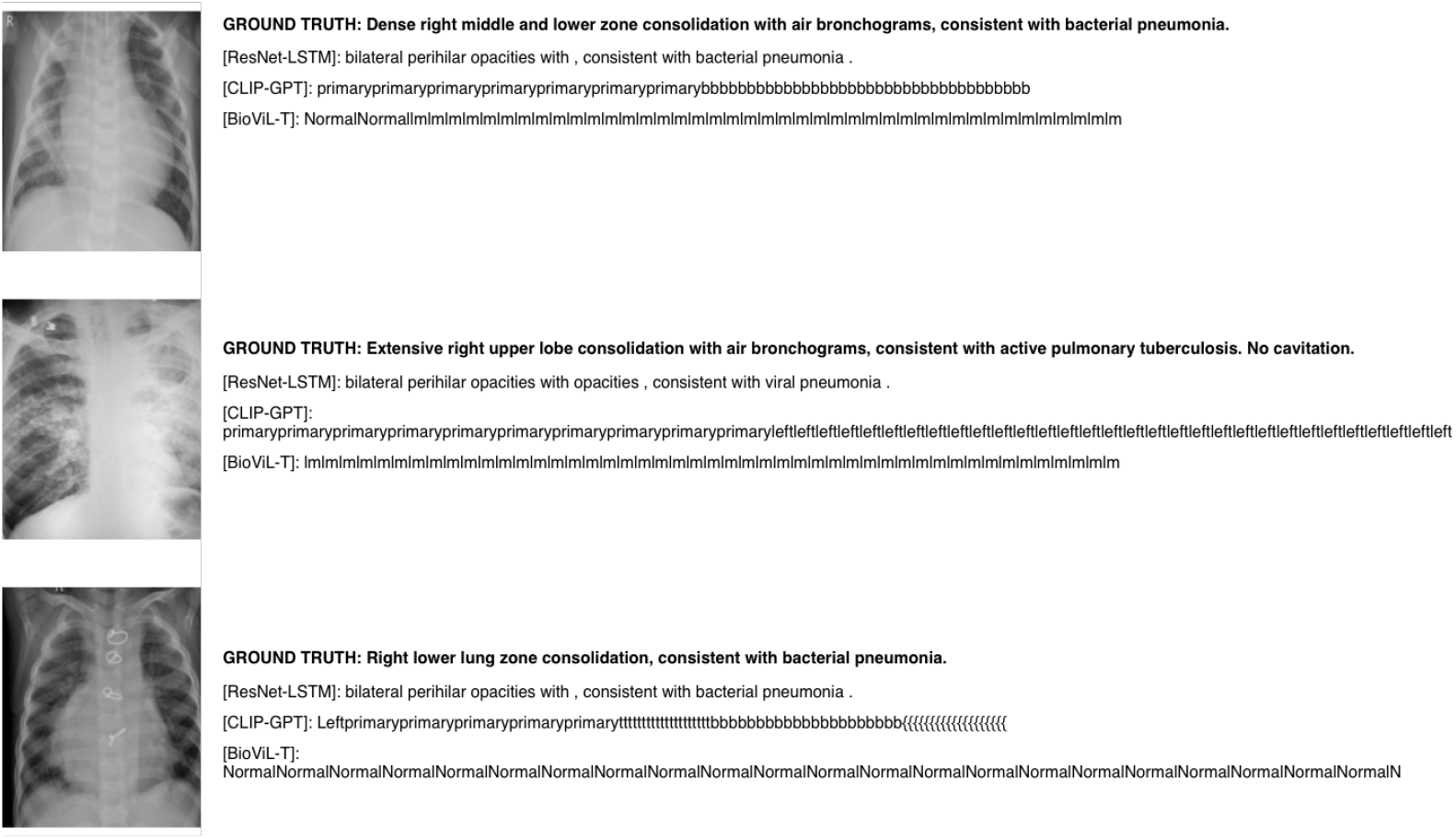
Extended qualitative comparison of generated reports. A larger set of generated reports, illustrating the specific failure modes of the V-L models (repetition loops vs. empty strings) compared with the ground truth.

**S2 Fig.**
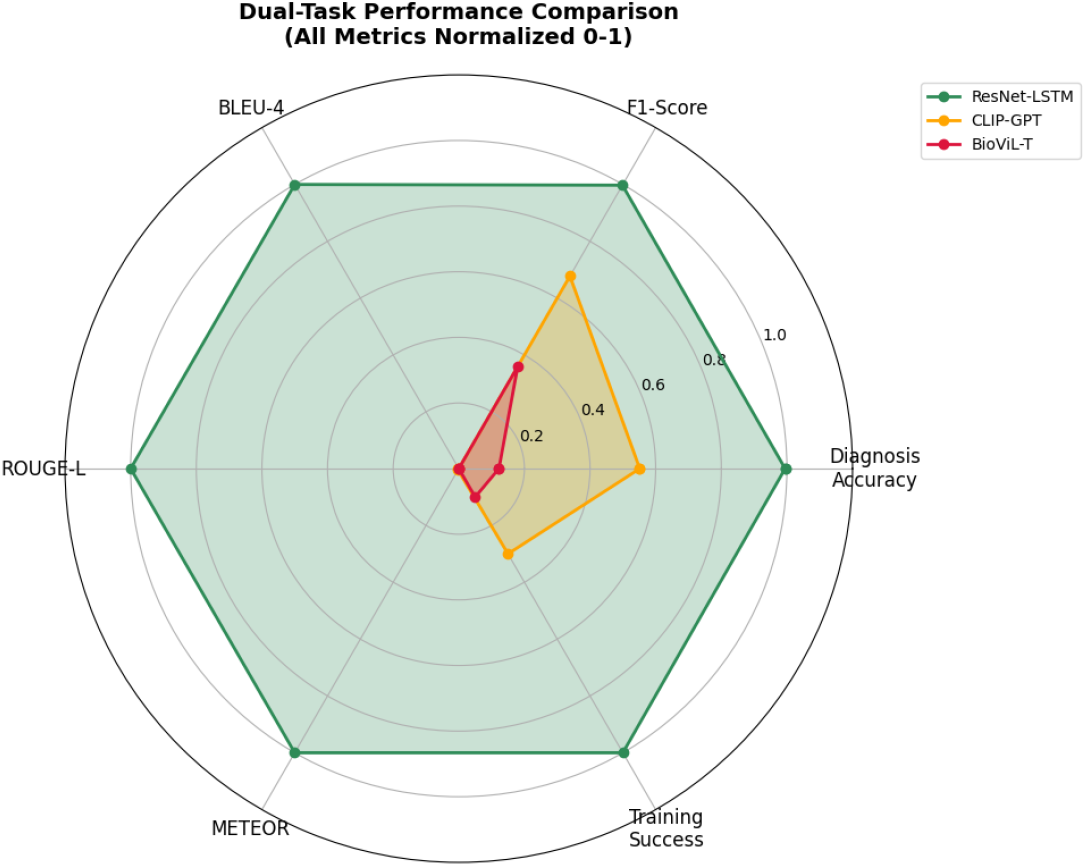
Holistic performance view. A normalized radar chart (0–1 scale) comparing the three architectures across six key metrics spanning classification, generation, and training stability. The ResNet-LSTM (green) maximizes the area across all diagnostic and generative metrics. In contrast, the modern V-L models (orange/red) show a collapsed profile, indicating that their failure is not specific to one metric but pervasive across all task dimensions.

**S3 Fig.**
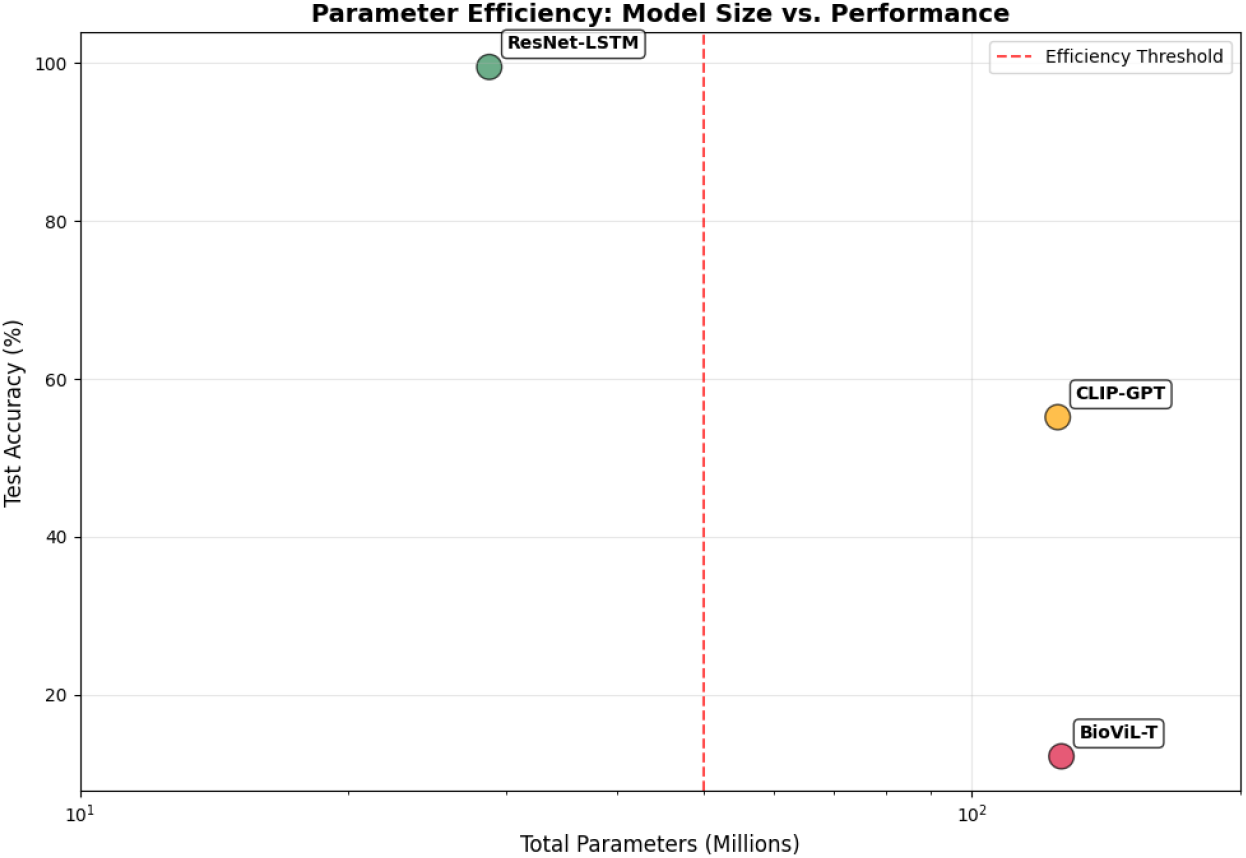
Parameter efficiency versus accuracy. The ResNet-LSTM (green) is both the smallest and the most accurate model; scale (120M+ parameters) does not correlate with performance on this specialized task.

## Acknowledgments

[TODO: Acknowledgments.]

